# Glucocorticoids regulate small extracellular vesicle (sEV) release via activation of nSMase2

**DOI:** 10.1101/2025.10.07.680983

**Authors:** Mia Burke, Clarissa Waites

## Abstract

Chronic stress, marked by prolonged elevation of glucocorticoid (GC) stress hormones, is a major risk factor for Alzheimer’s disease (AD) and accelerates AD pathology in mouse models. A key mechanism contributing to AD progression is the release of small extracellular vesicles (sEVs) carrying pathogenic proteins (e.g., tau, amyloid-beta) between brain regions, but the role of GCs in sEV biogenesis and release is unknown. Using total internal reflection fluorescence (TIRF) microscopy and the pH-sensitive marker mCh-CD63-pHluorin to visualize sEV release, we show that GCs stimulate sEV secretion in a neuronal cell line. This process requires the GTPase Rab27a and the enzyme neutral sphingomyelinase 2 (nSMase2), which catalyzes ceramide production and drives sEV formation. We further demonstrate that GCs promote sEV release by activating nSMase2 downstream of mitochondrial reactive oxygen species production and opening of the mitochondrial permeability transition pore (mPTP). These findings link GC-induced mitochondrial damage, specifically mPTP opening, to nSMase2 activation and enhanced sEV release by neuronal cells.

**Summary:** This study reveals a mechanism by which glucocorticoid stress hormones promote extracellular vesicle release from neuronal cells by activating the enzyme neutral sphingomyelinase 2. These findings provide a possible explanation for how stress accelerates the spread of Alzheimer’s pathology in the brain.

**Graphical Abstract:** 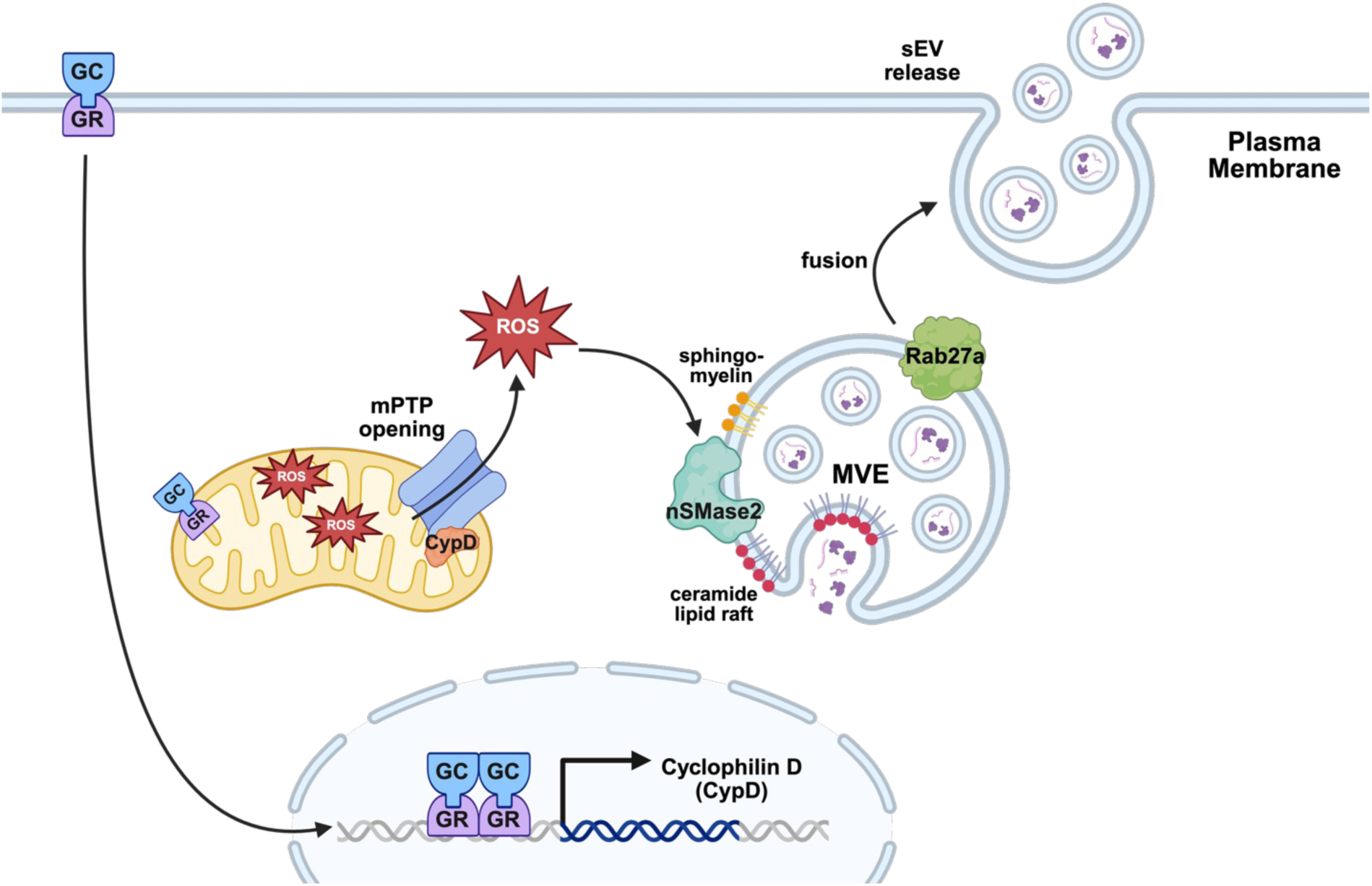

## Introduction

Chronic stress and the accompanying elevation of glucocorticoid (GC) stress hormones is among the strongest modifiable risk factors associated with Alzheimer’s disease (AD) [1–3]. Stress and GCs have been shown to initiate many pathogenic processes underlying AD progression, including the hyperphosphorylation and oligomerization of microtubule-associated tau protein and its secretion and propagation, the misprocessing of amyloid precursor protein into neurotoxic amyloid-beta (Aβ) peptides, and the activation of microglia and pro-inflammatory signaling pathways [4]. Furthermore, stress/GCs accelerate the onset of AD clinical symptoms and the spread of AD pathology through the brain, although the mechanisms responsible for this spread are not well understood [5–8].

Our recent work demonstrates that GCs drive the transmission of pathogenic tau through the hippocampus by stimulating its secretion via the type I unconventional secretory pathway. In this form of vesicle-free secretion, phosphorylated tau is translocated across the plasma membrane through interactions with specific lipids and heparin sulfate proteoglycans [7]. However, vesicle-mediated mechanisms are also implicated in the secretion and spread of AD-associated proteins and signaling molecules [9–11]. In particular, extracellular vesicles (EVs) have been shown to carry hyperphosphorylated tau, Aβ peptides and oligomers, inflammatory cytokines, and small noncoding/regulatory RNAs [12–16]. Of these, small EVs (sEVs), a defined subset of EVs with 50-150 nm diameter that originate in the endosomal pathway [17, 18], have been extensively characterized in AD and other neurodegenerative diseases. Not only are sEVs present in a range of accessible biofluids, including blood, cerebrospinal fluid, urine, and saliva [19–22], but sEV profiling has been used to successfully predict AD >10 years prior to the appearance of clinical symptoms [20]. These findings underscore the relevance of sEVs and their cargoes for AD diagnosis and progression, and the importance of understanding how their biogenesis and secretion are regulated by GCs and other stimuli.

The secretion of sEVs from cells occurs following fusion of multi-vesicular endosomes (MVEs) with the plasma membrane [17]. During fusion, the intraluminal vesicles (ILVs) formed through inward budding of the limiting membrane of the MVE are released to the extracellular environment and thereafter considered sEVs. Two main cellular pathways responsible for ILV budding and cargo loading are the endosomal sorting complex required for transport (ESCRT) and the neutral sphingomyelinase 2 (nSMase2)/ ceramide synthesis pathway [17, 23, 24]. In the former, a series of protein complexes (ESCRT-0, I, II, III) capture cargo and catalyze ILV formation, while in the latter, nSMase2-mediated synthesis of the bioactive lipid ceramide in condensed lipid raft domains drives negative membrane curvature and ILV formation [23, 25, 26]. nSMase2 is a phospho-activated enzyme that synthesizes ceramide from sphingomyelin, and thus serves as a regulatory checkpoint for membrane ceramide composition [27–30]. Intriguingly, nSMase2 is linked to AD pathophysiology, as its expression is elevated in AD brain tissue [31, 32], and its inhibition has been found to decrease the spread of both amyloid and tau pathology in mouse models [11, 31, 33, 34]. Furthermore, nSMase2 activity increases in response to multiple intracellular and extracellular stimuli, including reactive oxygen species (ROS), inflammatory cytokines, and disease-associated molecular patterns such as Aβ peptides and extracellular ATP [27, 35–40]. GCs can induce all of these stimuli, indicating their potential ability to regulate nSMase2/ceramide-associated sEV biogenesis and the subsequent release and spreading of pathogenic molecules [4].

In the current study, we investigate the role of GCs in sEV biogenesis and secretion using total internal reflection fluorescence (TIRF) microscopy coupled with a pH-sensitive marker of sEV release. We find that GCs stimulate sEV release in the neuronal Neuro2a cell line, and that this process requires nSMase2 activation. Additionally, we show that GC-induced nSMase2 activation is triggered by accumulation of mitochondrial ROS and opening of the mitochondrial permeability transition pore (mPTP), implicating mitochondrial damage as a driver of sEV release during stress/GC exposure. These findings reveal a GC-regulated mechanism of cell-cell transmission of biomolecules via sEVs, potentially contributing to the spread of pathology in AD.

## Results

### High glucocorticoid levels induce sEV release

To determine whether GC exposure alters sEV release in neuronal cells, we transfected Neuro2a (N2a) cells, a neuroblastoma cell line, with an mCh-CD63-pHluorin construct. CD63 is a tetraspanin enriched on the membranes of sEVs derived from MVEs in the endolysosomal pathway, and the dual fluorescent tags enabled us to identify cells expressing this construct (via constitutive mCherry fluorescence) as well as sEV release events (via unquenching of the pH-sensitive pHluorin moiety upon exposure to the neutral extracellular pH; **Fig. 1A**). The appearance of pHluorin signal at the cell surface was detected by total internal reflection fluorescence (TIRF) microscopy. Prior to imaging, N2a cells were treated for 24 hours with vehicle (control) or dexamethasone (Dex), a synthetic glucocorticoid, at a concentration known to activate glucocorticoid receptors (GRs) and elicit the cellular effects of chronic stress (5 uM; [7, 41]) Release of sEVs was then measured by quantifying the number of CD63-pHluorin+ events, defined as punctate flashes between 2-6 pixels (0.32-0.96 um) in size with characteristic rapid increases in fluorescence intensity (**Fig. 1B, C; Supplemental Video 1**), during a 3-minute imaging period. Intriguingly, we found that the number of CD63-pHluorin+ events observed during imaging was significantly higher in cells treated with Dex compared to vehicle control (**Fig. 1D; Supplemental Video 2**). Immunoblotting showed that this effect was not due to a Dex-mediated increase in expression of our mCh-CD63-pHluorin construct (**Fig. S1A-C).** Moreover, the Dex-induced increase in CD63+ sEV release was blocked by co-application of the GR antagonist mifepristone (10uM; **Fig. 1D; Supplemental Video 3**), demonstrating the GR-dependence of these release events.

**Figure 1.**
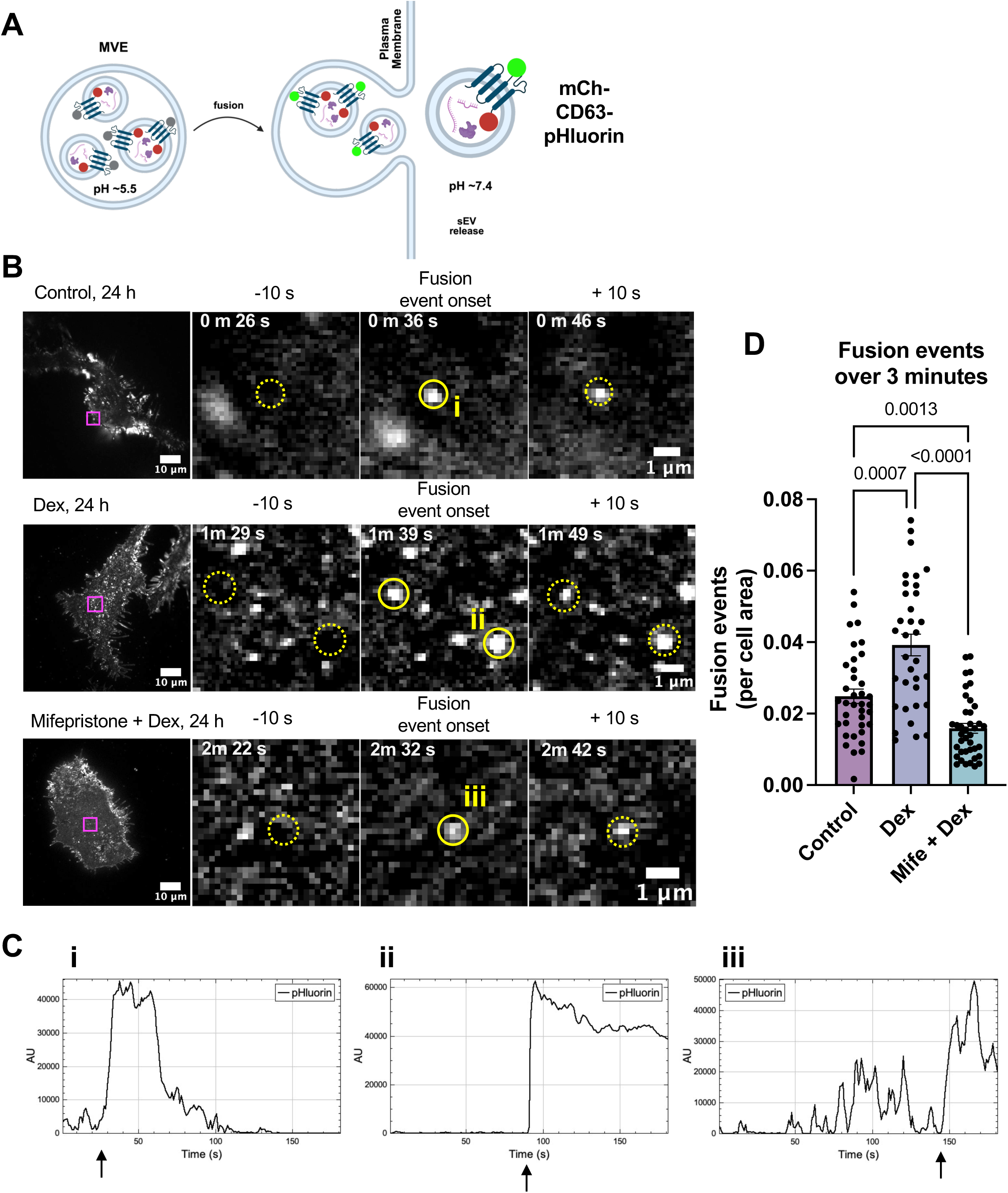
Glucocorticoids stimulate release of CD63+ sEVs. **A)** Schematic diagram showing how mCh-CD63-pHluorin is used to detect multivesicular endosome (MVE) fusion with the plasma membrane, based on appearance of green fluorescence upon exposure to the neutral extracellular pH. **B)** Maximum z-projection of pHluorin signal over 3 minutes of TIRF imaging in N2a cells treated for 24 h with vehicle (Control; top panel), 5uM dexamethasone (Dex; middle panel) or Dex + the GR antagonist mifepristone (10uM, Mife, bottom panel). Insets (magenta) are magnified to show the onset of individual fusion events (solid yellow circle), with frames depicting the same field of view 10 s before and after signal onset (yellow, dashed circle). **C)** Fusion events are characterized by the rapid onset of fluorescence (indicated by arrows), as shown in z-axis projections of fluorescence intensity over the 3 m imaging period. **i**, **ii**, and **iii** correspond to events identified for each condition in panel B. **D)** Quantification of fusion events, normalized to cell surface area (p values shown on graphs, Brown-Forsythe and Welch’s ANOVA with Dunnett’s correction for multiple comparisons; n=34-39 cells). Bars represent mean +/-SEM.

Since the presence of AD-relevant cargoes, including tau, have been well documented in sEVs isolated from both AD patients and animal models [11, 12, 20], we next examined whether tau could be detected in CD63+ sEVs. Here, we isolated sEVs from vehicle or Dex-treated N2a cells stably expressing human P301Ltau, a mutation associated with frontotemporal dementia and commonly used to study AD-related tauopathy [42, 43], and used the ExoView Imager to capture EVs and measure their size profiles and the presence of EV-enriched tetraspanins and oligomeric tau (**Fig. 2A**). Fluorescent immunolabeling with TOMA-1 and CD63 antibodies confirmed the colocalization of oligomeric tau with CD63+ sEVs isolated from both control and Dex-treated P301Ltau N2a cells (**Fig. 2B**). Interestingly, Dex treatment did not significantly change the percentage of TOMA+ sEVs (**Fig. 2C**), suggesting that GCs may enhance tau spreading by increasing overall sEV secretion rather than the fraction of tau+ sEVs. Dex treatment also did not alter the average size (**Fig. 2D**) or the tetraspanin composition of sEVs (**Fig. 2E, F**), suggesting that the effects we observe are due to changes in sEV biogenesis and release rather than to fundamental sEV characteristics.

**Figure 2.**
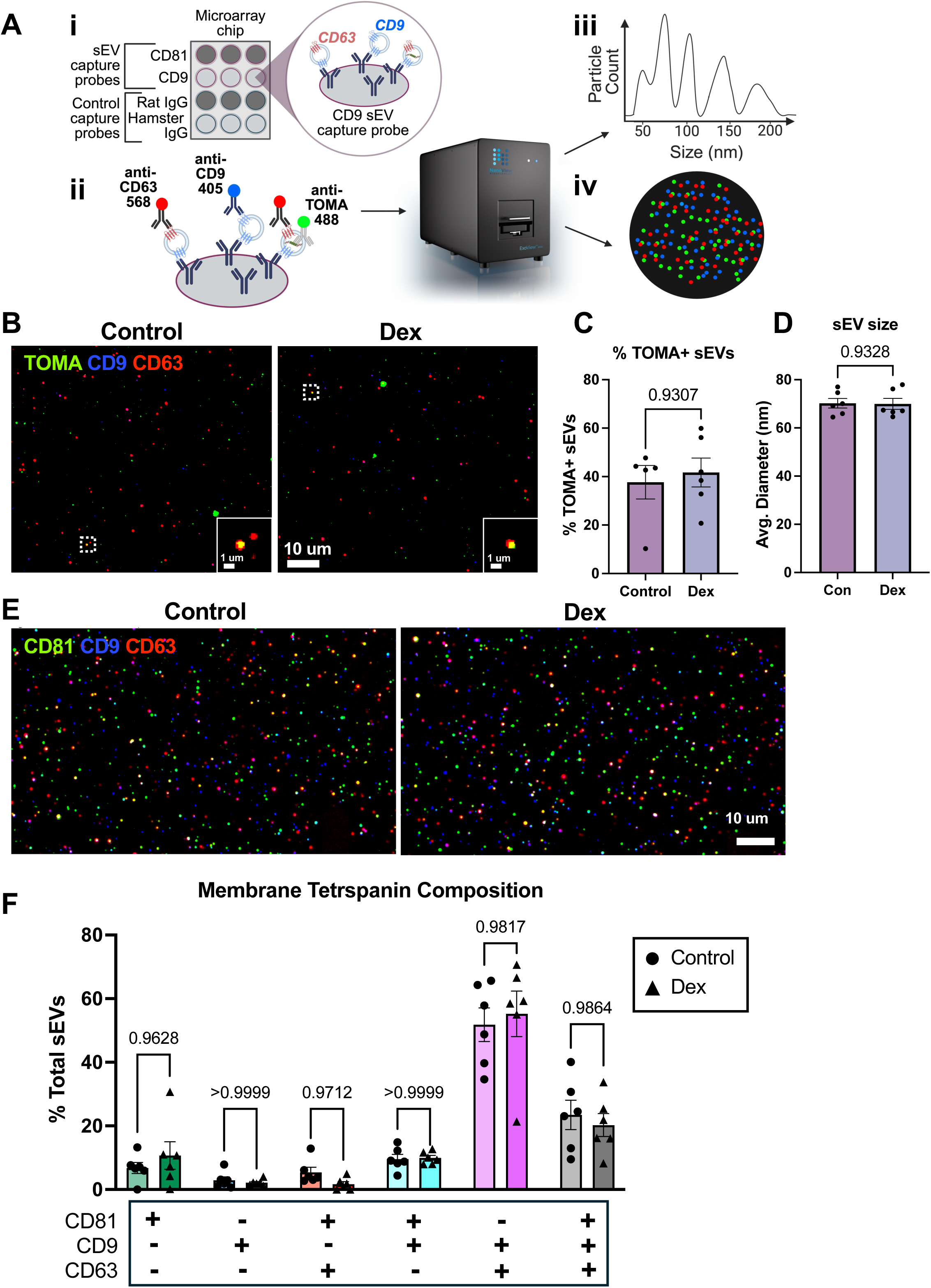
CD63+ sEVs can carry oligomeric tau. **A)** Schematic diagram of ExoView workflow, in which EVs are loaded onto capture chips coated with antibodies against EV-enriched membrane tetraspanins (CD81 and CD9) or control IgGs **(i)**, fluorescent antibodies are added to detect proteins on captured EVs **(ii)**, EV size distribution and number are measured by infrared interferometry **(iii)**, and EV protein composition is measured by immunofluorescence imaging **(iv)**. **B)** Representative images of sEVs isolated from P301Ltau N2a cells treated for 24 h with vehicle or Dex, fluorescently immunolabeled with TOMA-1 antibodies to detect intraluminal oligomeric tau (green), and antibodies against CD9 (blue) and CD63 (red) to detect sEV membrane-associated tetraspanins. Area indicated by white dotted box denotes the zoomed inset on the bottom left of each image, showing vesicles co-labelled for CD63 and TOMA. Scale bars as shown. **C)** Percent of TOMA+ sEVs from P301Ltau cells treated for 24 h with vehicle or Dex (p values on graph, Mann-Whitney U test, n=3 technical replicates for 5-6 biological replicates, bars represent mean +/-SEM). **D)** Average particle size of sEVs from P301Ltau cells treated for 24h with vehicle or Dex, measured by interferometric imaging. Raw size measurements were collated from technical replicates of both anti-CD81 and anti-CD9 spots for each sample (p values on graph, unpaired t-test, n=6, bars represent mean +/-SEM). **E)** Representative images of sEVs isolated from N2a cells treated with vehicle or Dex, fluorescently immunolabeled with CD81 (green), CD9 (blue), and CD63 (red) to detect sEV membrane tetraspanins. Scale bars as shown. **F)** Composition of CD81, CD63, and CD9 surface membrane tetraspanins found on P301LTau N2a-derived sEVs treated for 24 h with vehicle or Dex. Data are shown as percent of total sEVs in each sample that contain one, two, or all three tetraspanin(s), as indicated below the chart as “+” or “–” for each tetraspanin (p values on graph, 2-way ANOVA with Sidak’s multiple comparisons test, n=3 technical replicates for 5-6 biological replicates, bars represent mean +/-SEM). For all experiments, sEVs were purified by size exclusion chromatography (SEC) prior to overnight immobilization on anti-CD81 and anti-CD9 coated spots on microarray chips.

### Glucocorticoid-induced sEV release requires the small GTPase Rab27a

TIRF imaging of CD63-pHluorin has been used in multiple studies to visualize the release of MVE-derived sEVs [44–48], and we sought to confirm that we were imaging the same type of release event. Since the small GTPase Rab27a was previously shown to mediate MVE fusion with the plasma membrane resulting in sEV secretion [47, 49, 50], we examined CD63-pHluorin+ events following knockdown of Rab27a. Indeed, transfection of N2a cells with siRNAs against Rab27a led to a >40% reduction in Rab27a protein levels (**Fig. S1C, D**) and a >80% reduction in the number of CD63-pHluorin+ events/cell area induced by Dex compared to transfection with scrambled control siRNAs (**Fig. 3A; Supplemental Video 4**). To further confirm this finding, we assessed the colocalization of Rab27a with CD63-pHluorin+ events. When we co-expressed CD63-pHluorin with mCh-Rab27a in N2a cells and performed TIRF imaging, we found that these markers were often colocalized (**Fig. 3B**) and that mCh-Rab27a fluorescence peaked at the same time as that of CD63-pHluorin, although the onset of pHluorin signal occurred slightly ahead of the Rab27a signal (**Fig. 3C, D; Supplemental Video 5**). This lag is likely due to CD63-pHluorin+ sEVs being released at the time of MVE fusion, while Rab27a present on the limiting membrane of the MVE is not yet illuminated in the TIRF field. This high level of spatial and temporal colocalization with CD63-pHluorin was not observed with the late endosome/lysosome marker LAMP1 (**Fig. 3E, F; Supplemental Video 6**) or the autophagosome marker LC3 (**Fig. 3G, H; Supplemental Video 7**), indicating that CD63-pHluorin+ release events do not represent secretory lysosomes or autophagosomes, but rather sEVs released during MVE fusion with the plasma membrane.

**Figure 3.**
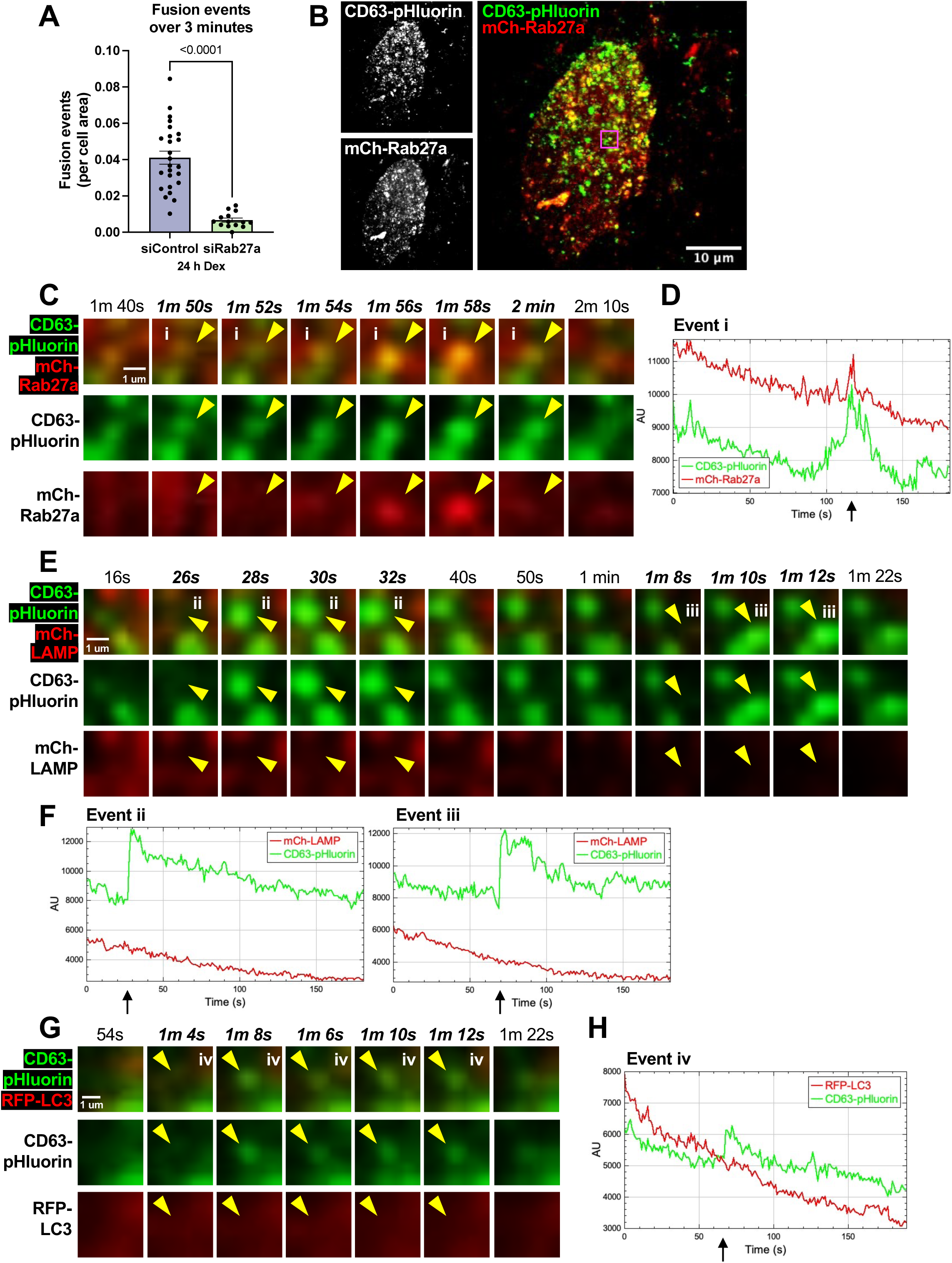
Rab27a is required for GC-induced sEV secretion and colocalizes with CD63+ fusion events. **A)** Quantification of CD63+ fusion events identified over 3-minute imaging periods, normalized to cell surface area. Dex-induced fusion events are significantly reduced by knockdown of Rab27a (p values on graph; unpaired t-test with Welch’s correction; n=14-25 cells). Bars represent the mean +/-SEM. **B)** Maximum z-projections of CD63-pHluorin and mCh-Rab27a over 3 minutes, with individual channels shown in greyscale and 2-color max projection in green (CD63-pHluorin) and red (mCh-Rab27a). The indicated area (magenta) corresponds to a single fusion event where both CD63-pHluorin and mCh-Rab27a are present. **C)** Time course of CD63-pHluorin and mCh-Rab27a fluorescence from the indicated area in panel B. Panels are shown at 2-second intervals over the duration of a single fusion event (yellow arrow) beginning 1 minute 50 seconds into the 3-minute video. **D)** Z-axis projection of CD63-pHluorin and mCh-Rab27a fluorescence intensities at the fusion event (**i**) indicated in panels B and C. Arrow indicates the onset of fusion/ CD63-pHluorin signal at 1 minute 52 seconds. **E)** Time course of CD63-pHluorin and mCh-LAMP1 fluorescence, magnified to show two individual fusion events. Panels are shown at 2 second intervals over the duration of two fusion events beginning 28 seconds (**ii**) and 1 minute 10 seconds (**iii**) into the 3-minute video, with the fusion events indicated by yellow arrows. Remaining panels are shown at 10 second intervals between onset of individual fusion events. **F)** Z-axis projection of CD63-pHluorin and mCh-LAMP1 fluorescence intensities over the 3-minute imaging time course at two fusion events (i**i** and **iii**), corresponding to the events in panel E. Arrows indicate the onset of fusion/ CD63-pHluorin signal. **G)** Time course of CD63-pHluorin and RFP-LC3 fluorescence, magnified to show individual fusion events. Panels are shown at 2 second intervals over the duration of the event beginning 1 minute, 6 seconds (**iv**) into the 3-minute video, with events indicated by yellow arrows. **H)** Z-axis projection of CD63-pHluorin and RFP-LC3 fluorescence intensities over the 3-minute imaging time course at an individual fusion event **(iv**), corresponding to events in panel G. Arrow indicates the onset of fusion/ CD63-pHluorin signal.

### The nSMase2/ceramide synthesis pathway drives GC-induced sEV release

To further probe the mechanism of GC-induced sEV release, we sought to identify the biogenesis machinery used to generate the GC-dependent subset of sEVs. One pathway known to mediate sEV biogenesis is the neutral sphingomyelinase 2 (nSMase2)/ceramide synthesis pathway responsible for hydrolyzing sphingomyelin into the bioactive lipid ceramide [23]. When ceramide accumulates in lipid raft domains, the conical shape of it head group generates sufficient negative membrane curvature to drive inward budding of the membrane and create intraluminal vesicles (ILVs) within the MVE [23, 25, 26]. Interestingly, nSMase2 is regulated by multiple stimuli (e.g., TNFα, Aβ plaques, oxidative stress) and like glucocorticoids, has been implicated in Alzheimer’s disease pathogenesis [27, 30, 35, 38]. To determine whether GC-driven sEV release occurs through the nSMase2/ceramide pathway, we again transfected N2a cells for 72 h with siRNAs to knockdown nSMase2 (gene name *Smpd3*) expression. Although we were unable to detect specific endogenous nSMase2 signal by immunoblotting with the available commercial antibodies, quantitative RT-PCR showed that siRNAs against nSMase2/Smpd3 (siSmpd3) led to a >75% reduction of mRNA (**Fig. S1E**). By TIRF imaging, we found that siSmpd3 significantly attenuated the Dex-induced increase in CD63-pHluorin+ sEV release compared to scrambled control siRNAs (**Fig. 4A-C; Supplemental Videos 8 and 9**). However, nSMase2 knockdown did not significantly alter CD63+ sEV release under baseline/control conditions (**Fig. 4C**), indicating that GC-induced sEV release in particular requires the nSMase2/ceramide synthesis pathway.

**Figure 4.**
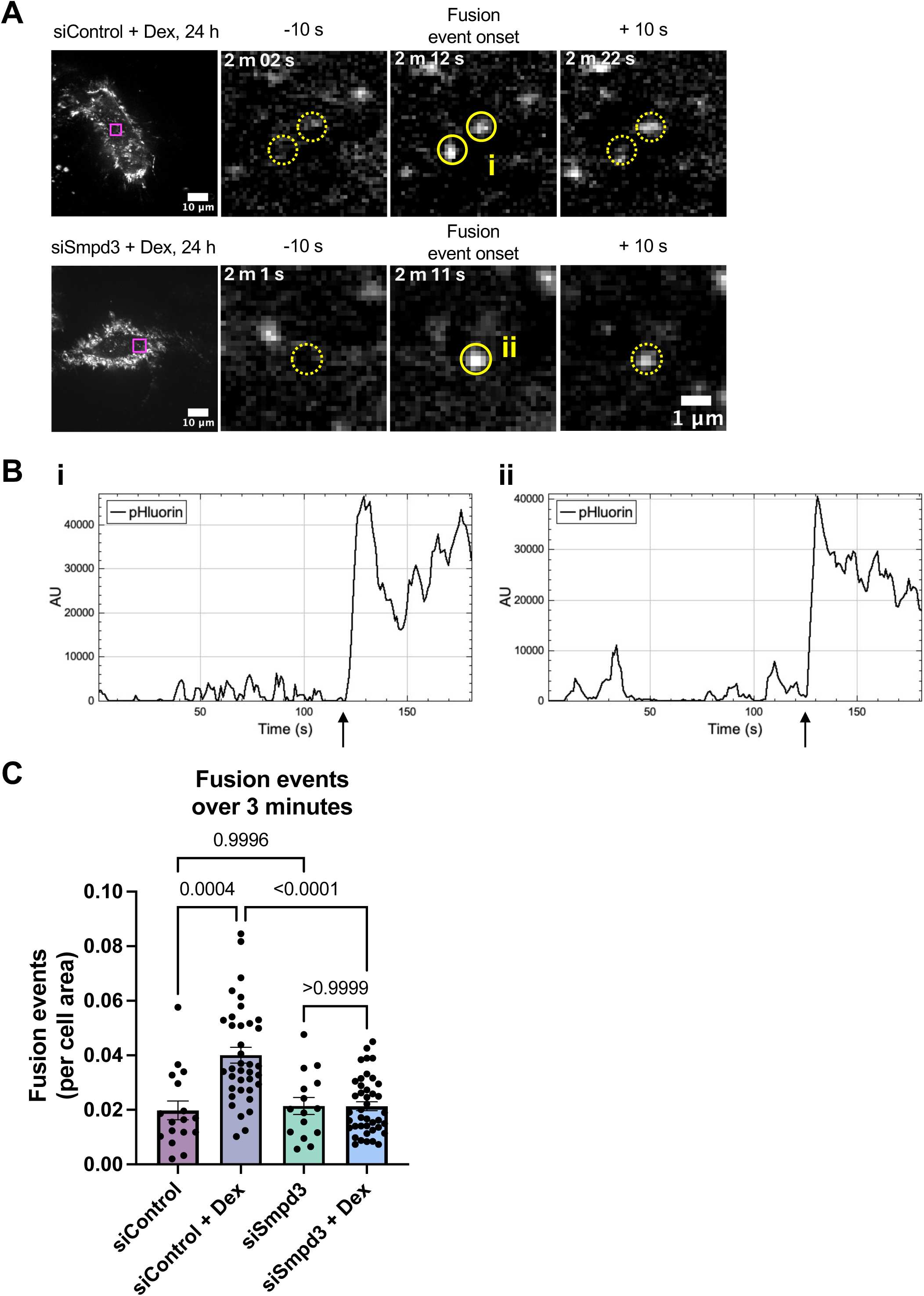
GC-dependent sEV release requires nSMase2. **A)** Maximum z-projection of pHluorin signal over the 3 m imaging period for siRNA control (top panel) and siSmpd3 (bottom panel) conditions. Insets (magenta) are magnified to show the onset of individual fusion events (solid yellow circles), with frames depicting the same field of view 10 s before and after signal onset (dashed yellow circles). **B)** Z-axis projections of pHluorin fluorescence intensity over the 3 m imaging period at events indicated by yellow circles for siControl (**i**) or siSmpd3 (**ii**). Arrows indicate the onset of an event/CD63-pHluorin signal, beginning at 59 s (**i**) or 1 minute, 31 seconds (**ii**). **C)** Quantification of fusion events identified over a 3 m imaging time course, normalized to the surface area of the cell (p values on graph, unpaired t-test with Welch’s correction, n=37-41 cells for Dex conditions and n=15-17 cells for vehicle conditions. For all graphs, bars represent mean +/-SEM.

### GCs increase nSMase2 activity and colocalization with sEV biogenesis machinery

Long-term effects of GCs typically occur through gene expression changes, as GC-bound GRs translocate to the nucleus and act as transcription factors [4, 51]. To determine whether GCs augment sEV biogenesis and release through transcriptional upregulation of nSMase2, we used quantitative RT-PCR to measure nSMase2/*Smpd3* mRNA levels in N2a cells treated with either vehicle or Dex for 24 hours. Here we found no significant differences, suggesting that GCs do not alter *Smpd3* expression at the transcriptional level (**Fig. 5A**). Since nSMase2 enzymatic activity is augmented by multiple intra- and extracellular stimuli, we evaluated whether it is similarly boosted by GC exposure. Using lysates of N2a cells over-expressing GFP-nSMase2 and treated with vehicle or Dex, we performed an enzyme-coupled fluorometric assay wherein nSMase2 activity is indirectly measured through resorufin fluorescence intensity. We found that our Dex treatment paradigm significantly increased nSMase2 activity (**Fig. 5B**). We also performed this assay in cell lysates following nSMase2 knockdown by siSmpd3. Due to the highly efficient nSMase2 knockdown, this experiment effectively gave us the level of background resorufin fluorescence due to non-specific sphingomyelinase activity, which has been subtracted from these data (**Fig. 5B**). Together, these findings indicate that GCs do not transcriptionally upregulate nSMase2/*Smpd3* but do significantly enhance its enzymatic activity.

**Figure 5.**
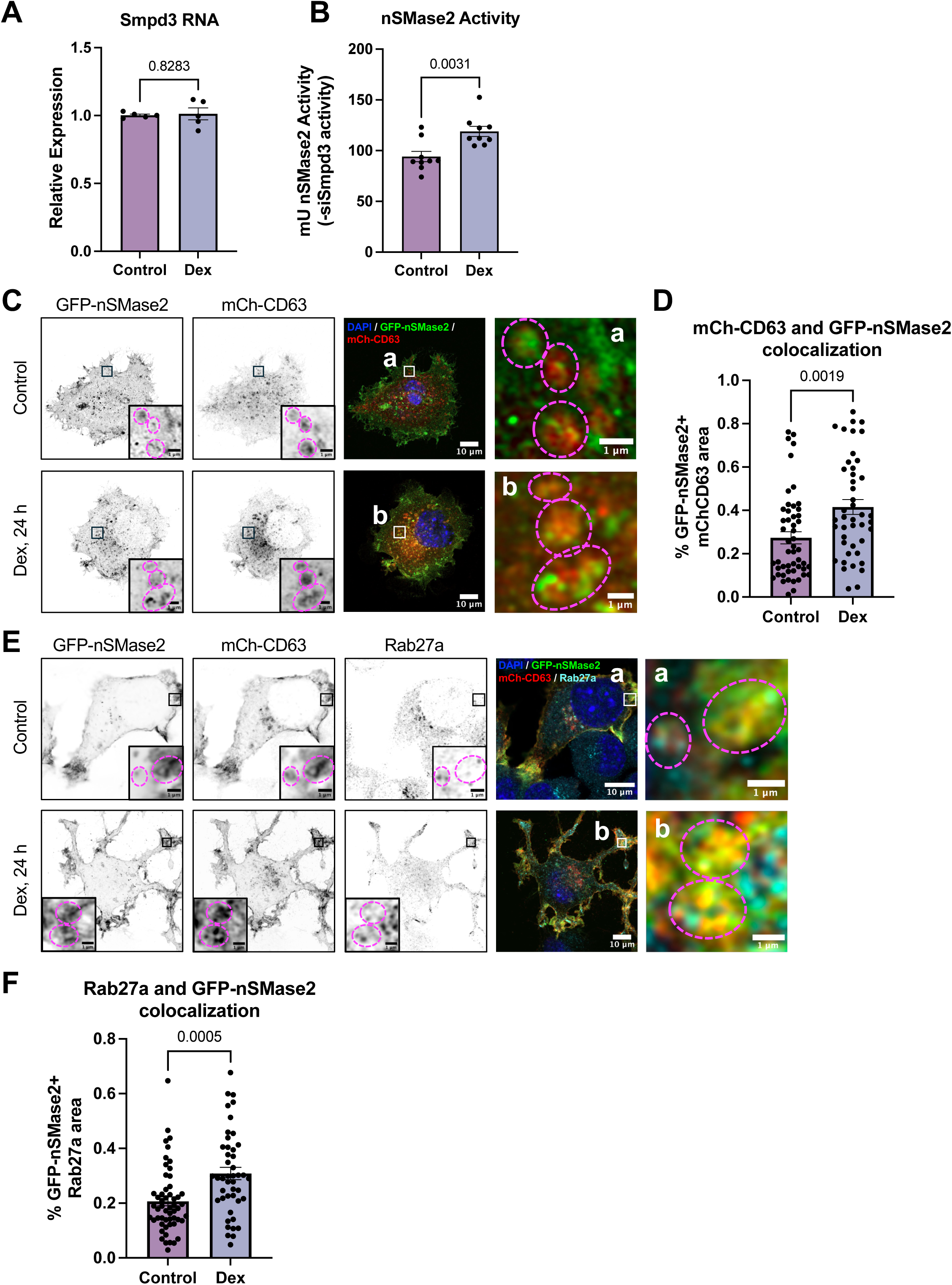
GCs increase nSMase2 activation and colocalization with sEV biogenesis machinery. **A)** Quantification of mRNA levels of nSMase2/Smpd3 24 h after treatment with Dex or vehicle control (p values on graph, unpaired t-test with Welch’s correction, n=5). **B)** Quantification of nSMase2 activity 24 h after treatment with Dex or vehicle control. nSMase2-specific activity was determined by subtracting the mean activity observed in cells with >75% knockdown of nSMase2 by siSmpd3 (p values on graph, unpaired t-test with Welch’s correction, n=9). **C)** Super-resolution images of N2a cells expressing GFP-nSMase2 (green) and mCh-CD63 (red), stained for DAPI (blue) and treated with vehicle or Dex for 24 h. Zoomed insets for control (**a**) and Dex (**b**) conditions, indicated by black squares in black/white images and white squares in color images, are shown in color (far right panels). Dashed magenta circles indicate the areas of colocalization for GFP-nSMase2 and mCh-CD63. **D)** Quantification of mCh-CD63 and GFP-nSMase2 colocalization, represented as the percent of mCh-CD63 area that is GFP-nSMase2+ as determined by Mander’s colocalization coefficient (p values on graph, unpaired t-test with Welch’s correction, n=44-51 cells). **E)** Super-resolution images of N2a cells expressing GFP-nSMase2 (green) and mCh-CD63 (red), stained for DAPI (blue) and Rab27a (cyan) and treated with either vehicle or Dex for 24 h. Zoomed insets for control (**a**) and Dex (**b**) conditions, indicated by black squares in black/white images and white squares in color images, are shown in color (far right panels). Dashed magenta circles indicate the areas of colocalization for GFP-nSMase2, mCh-CD63, and Rab27a. **F)** Quantification of Rab27a and GFP-nSMase2 colocalization, represented as the percent of Rab27a area that is GFP-nSMase2+ as determined by Mander’s colocalization coefficient (p values on graph, unpaired t-test with Welch’s correction, n=45-54 cells). For all graphs, bars represent mean +/-SEM.

To determine whether increased nSMase2 enzymatic activity corresponds to changes in its subcellular localization, we measured nSMase2 colocalization with CD63 and Rab27a, both markers of MVEs/sEVs. Using super-resolution imaging (Zeiss Airyscan), we first assessed whether 24-hour Dex treatment stimulated the association of GFP-nSMase2 with mCh-CD63 in N2a cells. While we observed instances of GFP-nSMase2 and mCh-CD63 colocalization in both vehicle and Dex treatment conditions, Mander’s colocalization analysis revealed that Dex significantly increased the percentage of mCh-CD63+ structures that were also positive for GFP-nSMase2 (**Fig. 5C, D**). Immunostaining with antibodies against Rab27a further revealed that these GFP-nSMase2+/mCh-CD63+ structures colocalize with endogenous Rab27a, and that Dex increases the association of GFP-nSMase2 with Rab27a (**Fig. 5E, F**). These findings are in agreement with a previous report of increased association between nSMase2 and Rab27a following TNFα-mediated nSMase2 activation [28]. We also assessed the localization of GFP-nSMase2 to early endosomes and late endosomes/MVEs, marked by immunostaining with antibodies against EEA1 and Rab7, respectively. These experiments revealed the presence of GFP-nSMase2 in both compartments under control and Dex treatment conditions (**Fig. S2**), indicating that nSmase2 is found throughout the endosomal pathway, where sEV biogenesis occurs. While Dex treatment did not elicit a change in GFP-nSMase2 colocalization with EEA1+ early endosomes (**Fig. S2A, C**), increased colocalization was observed with Rab7+ late endosomes (**Fig. S2B, D**). These findings support the concept that GC-mediated nSMase2 activation leads to its increased association with late endosomes/MVEs, where it can locally produce ceramide to catalyze sEV biogenesis and secretion.

### GC-induced mROS production and mPTP opening stimulate nSMase2 activation and sEV release

Previous studies have shown that nSMase2 activity can be regulated by mitochondrial reactive oxygen species (mROS)-mediated signaling cascades. For instance, PKC and MAPK activation are triggered by mROS-mediated oxidation [51–54] and implicated in nSMase2 phospho-activation [40]. Additionally, mROS are known to inhibit the phosphatase calcineurin, which itself is an inhibitor of nSMase2 activity [40]. We recently demonstrated that chronic GC exposure increases cytosolic mROS levels by stimulating opening of the mitochondrial permeability transition pore (mPTP) via transcriptional upregulation of the mPTP activating component cyclophilin D [55]. Based on these findings, we assessed whether mROS and mPTP opening are required for the increase in GC-induced sEV release. First, we confirmed that our GC treatment paradigm induced mROS production in N2a cells by measuring the fluorescence intensity of HyPerRed-Mito, a genetically encoded reporter of mROS [56]. As anticipated, 24 h Dex treatment led to a ∼50% increase in HyPerRed-Mito intensity compared to vehicle (**Fig 6A, B**). We next monitored GC-induced mCh-CD63-pHluorin sEV release during pharmacological inhibition of mROS and mPTP opening (**Fig. 6C**). Here, N2a cells were treated for 24 h with Dex and either vehicle, MitoTEMPO (a mitochondrially-targeted ROS scavenger that mimics superoxide dismutase [57]), cyclosporin A (CysA; a cyclophilin D inhibitor that blocks mPTP opening [58]), or mito-apocynin (mitoApo; a mitochondrially-targeted inhibitor of the ROS producing enzyme NADPH oxidase [59]). TIRF imaging revealed that each of these inhibitors effectively block the Dex-induced increase in CD63+ sEV release (**Fig. 6D; Supplemental Video 10**), indicating that mROS and mPTP opening are drivers of GC-mediated sEV release. This finding supports previous work identifying ROS as a driver of EV release in PC12 pheochromocytoma and M1 renal epithelial cells [60–63]. Moreover, using the fluorometric nSMase2 activity assay, we found that these inhibitors significantly decreased Dex-induced nSMase2 activation (**Fig. 6E**). Together, our data suggest a mechanism wherein GCs induce mPTP opening and mROS production, leading to nSMase2 activation, which in turn drives sEV biogenesis and release.

**Figure 6.**
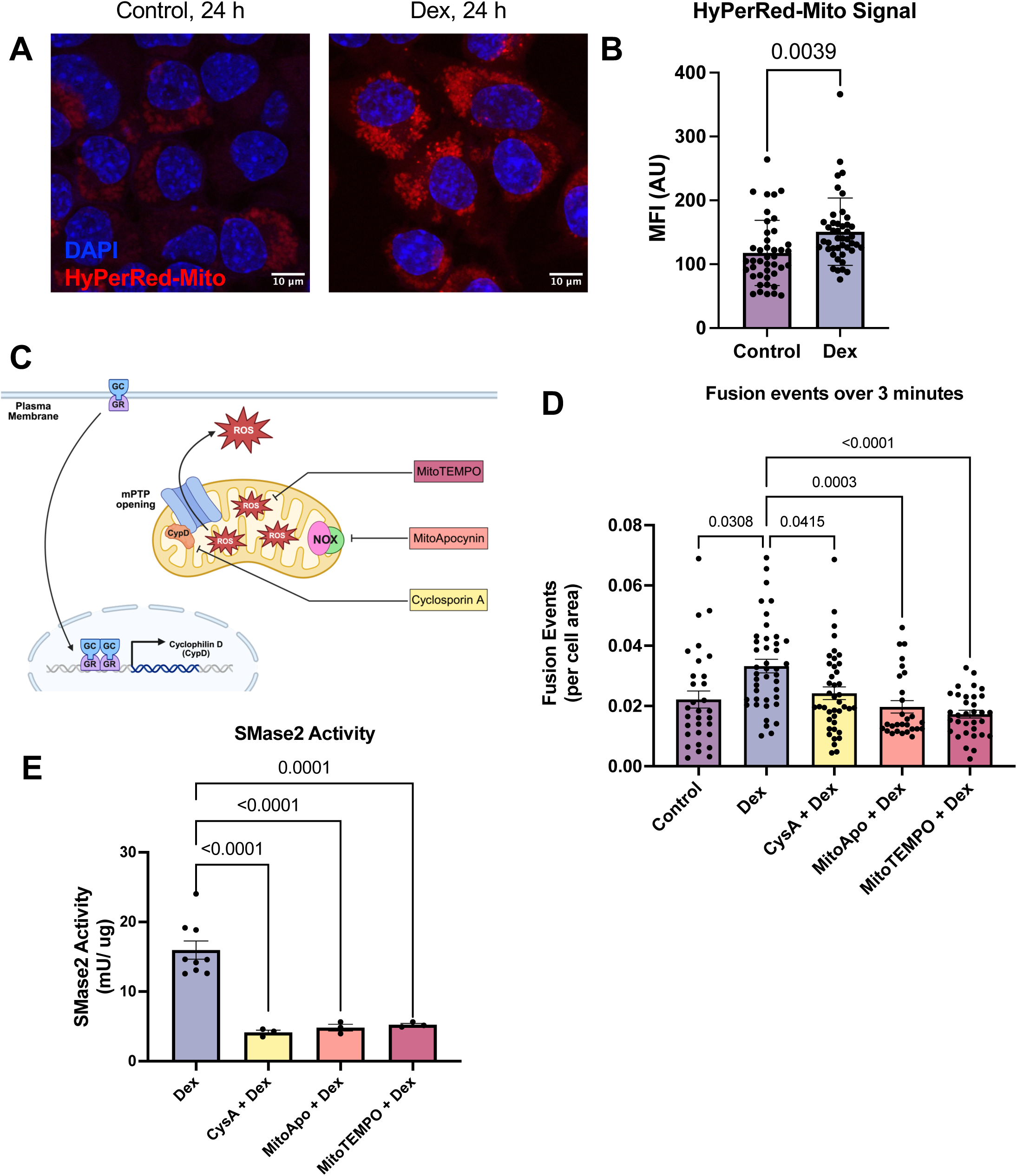
GCs drive sEV release through mPTP opening and mROS accumulation. **A)** Representative images of N2a cells expressing HyPerRed-Mito (red) and stained with DAPI (blue) following treatment with vehicle (control) or Dex for 24 h. **B)** Quantification of mean fluorescence intensity (MFI) of HyPerRed-Mito in N2a cells treated with vehicle or Dex for 24 h (p values indicated, unpaired t-test with Welch’s correction, n=42-44 cells). **C)** Schematic depicting the targets of mitochondrial ROS inhibitors used in panels **D** and **E**. Created using BioRender. **D)** Quantification of fusion events identified over a 3 m imaging period, normalized to the cell surface area. Images were acquired after 24 h treatment with vehicle, Dex, or Dex + cyclosporin A (CysA), mito-apocynin (MitoApo), or mito-TEMPO (p values on graph, Brown-Forsythe and Welch’s ANOVA with Dunnett’s post-hoc analysis; n=28-42 cells). **E)** nSMase2 activity after 24 h treatment with Dex or Dex + CysA, MitoApo, or Mito-TEMPO. Activity was determined by enzyme-coupled fluorescent assay and normalized to ug total protein (p values on graph, Brown-Forsythe and Welch’s ANOVA with Dunnett’s post-hoc analysis, n= 3-9 replicate wells). For all graphs, bars represent mean +/-SEM.

## Discussion

In the current study, we show that GC stress hormones stimulate sEV secretion, and that these events are dependent upon the GTPase Rab27a, the ceramide synthesis enzyme nSMase2, and the accumulation of mROS. We first confirm that CD63-pHlourin+ events represent sEV release/fusion of MVEs with the plasma membrane by showing their dependence on the sEV-linked small GTPase Rab27a, and their colocalization with Rab27a but not autophagy or lysosomal proteins. We next show that GC-induced sEV release is mediated by nSMase2, as nSMase2 knockdown abolishes GC-mediated CD63+ events, while GC exposure promotes nSMase2 activation and localization to CD63+ and Rab27a+ structures. Finally, we demonstrate that GC-driven nSMase2 activation and CD63+ sEV release are blocked by pharmacological inhibitors of mROS and mPTP opening, indicating their dependence on high mROS levels. These experiments reveal a GC-induced mechanism driving sEV release and potentially sEV-mediated spreading of AD pathology, given our finding of oligomeric tau in CD63+ sEVs and many other studies reporting that sEVs carry AD-relevant cargoes [64, 65].

Our work clearly demonstrates that high GC levels drive sEV release, in agreement with a previous study showing that the chronic unpredictable stress paradigm elicits sEV release from hippocampal tissue [66]. To our knowledge, no other molecules produced by the HPA axis during stress (e.g., corticotropin releasing hormone, mineralocorticoids, etc.) have been implicated in sEV release. However, meta-analysis of more than 100 studies of >5,000 patients with major depressive disorder (MDD), closely linked to chronic stress, indicate elevation of several inflammatory cytokines (e.g., IL-1β, IL-2, IL-6, IL-12, and TNF-α) in the periphery [67]. Thus, it is possible that sEV release is driven in part by the effects of pro-inflammatory cytokines on biogenesis machinery, of which nSMase2 may serve as a central target given its known activation by TNFα and IL-1β [27]. Indeed, the combinatorial effect of GCs and pro-inflammatory cytokines, particularly when GC release precedes an inflammatory event, have been shown to amplify inflammatory responses through activation of the NLRP3 inflammasome [68, 69]. Whether this additive effect also exists for sEV release remains an open question. In our model, GC-induced sEV release is mediated through activation of nSMase2; thus, we surmise that the increased presence of nSMase2-activating stimuli (e.g., pro-inflammatory cytokines, extracellular Aβ, etc.) in the setting of physiological stress or the neuroinflammatory landscape of AD further amplifies nSMase2/ceramide sEV biogenesis and release. Moreover, we predict that any stimuli serving to enhance mROS production and release from mitochondria (e.g., glutamate excitotoxicity, inflammation, misfolded proteins or aggregates) upstream of nSMase2 activation will also augment sEV biogenesis and release [70].

CNS mitochondrial dysfunction has been well-characterized across many neurodegenerative diseases [71]. The brain accounts for nearly 20% of all energy consumption by the body, despite accounting for only 2% of the total body mass [72]. Loss of energetic homeostasis impairs the ability of neurons to carry out basic, energy-intensive functions, including maintenance of membrane potential using ion pumps and long-distance trafficking of key machinery between the soma and synapses [73–75]. Additionally, dysfunction of mitochondrial electron transport chain complexes can lead to build-up of ROS, which can leak from mitochondria and drive widespread cellular damage through redox of protein, DNA, and lipid targets [76]. The current study broadens these findings, demonstrating that mROS production is a driver of sEV-mediated cell-to-cell communication in the brain. Neuronal mROS-responsive sEV release may play a protective role, by shuttling excess biomolecules to nearby cells with greater degradative capacity (e.g., microglia), transmitting damage signals, or signaling the need for neurotrophic support. Under chronic GC exposure and associated mROS elevation, it is also possible that the role of sEV release changes over time, first acting as a neuroprotective mechanism and later conferring neurotoxicity as the primary dysfunction endures. Given the complexity of intercellular signaling during stress and neurodegeneration, further studies identifying the cells targeted by GC-induced sEVs and the composition of sEV cargoes over time will be necessary.

To date, little is known about the impact of GCs on sEV composition. While we identify oligomeric tau as a cargo of CD63+ sEVs secreted from N2a cells, we do not see a change in the ratio of tau+ sEVs following 24-hour exposure to GCs, nor do we see alterations in the membrane tetraspanins CD9, CD81, or CD63. Further investigation is needed to identify GC-enriched sEV proteins and biomolecules by unbiased screening methods; however, recent evidence suggests that GCs can regulate sEV cargo recruitment. In particular, the microRNA miR-335-5p is reportedly enriched in sEVs derived from muscle satellite cells treated with dexamethasone [77]. In neurons, miR-335-5p was found to target voltage-gated sodium channels and the glutamine receptor, GRM4 [78, 79], suggesting that GC-mediated enrichment of this miRNA in sEVs could facilitate the neuromodulatory effects of GCs. Additional data on how GCs may regulate sEV cargo comes from investigations of potential MDD biomarkers, which identify multiple miRNAs and proteins with roles across neurotransmission, metabolism, and inflammation [80, 81]; however, the biological relevance of these identified cargoes is unclear. In contrast to GCs, ROS have been directly implicated in sEV cargo composition. In particular, ROS-mediated redox of kinases and phosphatases was shown to regulate the recruitment of phosphorylated protein cargoes, including annexin A2 (ANXA2) and Ephrin A2 (EphA2), into nascent ILVs [60, 61, 82]. ANXA2-enriched sEVs play a protective role in recipient cells exposed to oxidative stress [61], and ANXA2 is associated with synapse formation, cell survival, and maintenance of blood-brain-barrier integrity in the nervous system. Thus, the potential enrichment of ANXA2 in sEVs whose biogenesis and release are triggered by elevated GCs and ROS may have a protective role on surrounding cells [83]. By contrast, EphA2-containing sEVs may have the opposite effect on CNS health by negatively regulating neurogenesis, amplifying neuroinflammation, and disrupting blood-brain-barrier integrity [84, 85]. Finally, nSMase2 itself appears to regulate sEV cargo composition, given its role in the spreading of tau and oligomeric Aβ in AD mouse models, and its ability to regulate the levels of several sEV-enriched microRNAs promoting angiogenesis and cell migration in cancer [11, 86–89]. Together, these findings suggest multiple potential points of cargo regulation in the GC-mediated pathway for sEV biogenesis, representing a rich area for future studies.

Chronic stress is a major risk factor for AD, and GC-induced effects on neuronal and glial function appear to hasten disease progression. If GC-driven sEV release contributes to the spread of pathogenic molecules across brain regions during the prodromal stage of AD, targeting this mechanism will be imperative to delaying or preventing disease progression. There are several potential therapeutic targets in the pathway we identify in this study [55, 90]. In particular, we previously showed that inhibition of mPTP opening and the mPTP activator CypD can attenuate GC-induced mitochondrial dysfunction and tau hyperphosphorylation and oligomerization in murine neurons. While clinical use of the CypD inhibitor cyclosporin A is untenable given its toxicity and inability to cross the blood-brain barrier, other CypD inhibitors are currently under development [91–94]. Moreover, we demonstrated the efficacy of the NAPDH oxidase inhibitor mito-apocynin in preventing GC-induced tau pathology, synapse loss and behavioral changes in the murine brain, and pharmacokinetic and toxicology studies of this compound in mice are promising [55, 95]. The superoxide dismutase mimetic mito-TEMPO was similarly shown to reduce tau oligomerization in AD mitochondrial cybrid cells and has been used to successfully treat mouse models of metastatic breast cancer, diabetic cardiomyopathy, and acute hepatotoxicity [96–98]; however, further investigation of mito-TEMPO’s bioavailability in the brain, as well as its efficacy against neurodegeneration, is needed [99, 100]. Downstream of mROS, nSMase2 is also an attractive druggable target for the treatment of neurodegenerative conditions. Indeed, nSMase2 inhibitors have shown promise in mitigating pathology in mouse models of AD [11, 33, 34], Parkinson’s disease, and other neuroinflammatory conditions including HIV-associated dementia [101], and many small molecule inhibitors have recently been described [90]. Finally, targeting Rab27a is another strategy that has been considered for attenuating the spread of neurodegenerative pathologies [102]; however, the complexity of targeting both Rab27a and Rab27b isoforms and the widespread roles of this GTPase throughout the body limits the utility of such a therapy. Recent identification of the Rab27a-stabilizing adaptor protein KIBRA, and its role in sEV release, could allow for the development of inhibitors targeting the specific interaction between Rab27a and KIBRA, such as was done for Rab27a and the adaptor protein JFC1 [103, 104].

In summary, this study has uncovered a key cellular pathway linking elevated GC levels to sEV biogenesis and release. Specifically, we show that mROS, produced in response to GC-mediated mitochondrial damage, promote the activation of nSMase2 and drive its association with relevant MVE biogenesis and fusion machinery, culminating in sEV release. Given the extensive evidence that pathogenic molecules are present in sEVs and contribute to disease progression during the course of AD and other neurodegenerative conditions, this work demonstrates a potential mechanism whereby elevated GCs accelerate AD development.

## List of Abbreviations

Aβ: amyloid beta
AD: Alzheimer’s disease
ANXA2: annexin A2
CysA: cyclosporin A
Dex: dexamethasone
EphA2: ephrin A2
ESCRT: endosomal sorting complexes required for transport
EVs: extracellular vesicles
GC: glucocorticoid
GR: glucocorticoid receptor
ILVs: intraluminal vesicles
MDD: Major Depressive Disorder
MitoApo: mito-apocynin
mPTP: mitochondrial permeability transition pore
mROS: mitochondrial reactive oxygen species
MVEs: multi-vesicular endosomes
N2a: Neuro-2a
nSMase2: neutral sphingomyelinase 2
ROS: reactive oxygen species
sEV: small extracellular vesicle
TIRF: total internal reflection fluorescence

## Materials and Methods

### Cell culture and cell lines

Neuro2a (N2a) neuroblastoma cells (ATCC, CCL-131) were cultured in DMEM/GlutaMAX (ThermoFisher Scientific/Invitrogen) with 10% FBS (Atlanta Biological) and Antibiotic–Antimycotic (ThermoFisher Scientific/Invitrogen) and kept at 37°C in 5% CO2. N2a cells stably transfected with P301LhTau-EGFP (2N4R) were a kind gift from Professor Juergen Gotz (University of Queensland, Australia) and Ioannis Sotiropoulos (NCSR Demokritos, Greece) and cultured in an identical manner to wildtype N2a. For all experiments, cells were used between 5 and 25 passages. For live-imaging experiments, N2a cells were plated onto glass-bottom 50-mm dishes (MatTek; 35,000-50,000 cells/dish) and transfected using Lipofectamine 3000 (ThermoFisher Scientific/Invitrogen) according to manufacturer’s instructions. Cells were imaged or fixed and immunostained 24-48 hours later. For siRNA experiments, N2a cells were transfected with 37.5 pmol (50-mm dish, 6-well plate per well) or 187.5 pmol (10-cm dish) silencer select siRNAs against nSMase2/Smpd3 (s81737, ThermoFisher Scientific/Invitrogen), Rab27a (s201063, Thermo Fisher Scientific/Invitrogen), or control siRNAs (4390843, ThermoFisher Scientific/Invitrogen) using Lipofectamine 2000 (ThermoFisher Scientific/Invitrogen) in culture media without antibiotic, according to the manufacturer’s instructions. Cells were imaged or harvested 72 h later. For sEV purification followed by ExoView EV-on-a-chip analyses, cells were initially cultured as usual, but media was later exchanged for HEK media prepared with 10 % exosome-depleted FBS (Gibco; Cat No. A2720803) at the time of cell treatment.

### DNA plasmids

pLenti-pHluorin_M153R-CD63 was described in Sung et al (2020) (Addgene #172117) [46]; pLAMP1-mCh was described in Van Engelenburg et al (2010) (Addgene #45147) [105]; pmRFP-LC3 was described in Kimura et al (2007) (Addgene #21075); pC1-HyPeRed-mito was described in Ermakova et al (2014) (Addgene #60247) [56]. Human CD63 (GenBank ID: CR542096.1) tagged with pHluorin_M153R in a small extracellular loop as described by Sung et al (2020) [46] was synthesized at Genewiz, subcloned into the pC2-mCherry vector (Clonetech) at the Kpn1 and BamH1 digestion sites. mCh-Rab27a was generated from digestion of pEGFP-Rab27a C1 (Westbroek et al 2008, Addgene #89237 [106]) where mCh derived from a mCh-C1 vector was inserted at the AgeI and XhoI sites. Mouse nSMase2 (GenBank ID: NM_021491.4) was synthesized (Genewiz) in pUC-GW-Kan-nSMase2 and cloned into the pEGFP-N1 vector backbone at the BamH1 digestion site.

### Antibodies and pharmacological agents

The following antibodies were used: tubulin alpha mouse monoclonal antibody (T9026; Sigma), GFP rabbit polyclonal antibody (A-6455; Invitrogen), mCh rabbit polyclonal antibody (PA5-34974; Invitrogen), Rab27a rabbit monoclonal antibody (D7Z9Q; Cell Signaling Technologies), EEA1 rabbit monoclonal antibody (C45B10; Cell Signaling Technology), Rab7 mouse monoclonal antibody (ab50533; Abcam), HGS/ Hrs rabbit polyclonal antibody (ab155539; Abcam), CHMP-4b rabbit polyclonal antibody (13683-1-AP; Proteintech), TOM20 rabbit monoclonal antibody (D8T4N; Cell Signaling Technology), Alexa Fluor 647 goat anti-rabbit IgG highly cross-adsorbed secondary antibody (A-21245; ThermoFisher Scientific/ Invitrogen), Alexa Fluor 647 goat anti-mouse IgG highly cross-adsorbed secondary antibody (A-21235; ThermoFisher Scientific/ Invitrogen), IRDye 800CW goat anti-mouse IgG secondary antibody (P/N: 926-32210, LI-COR), IRDye 680CW goat anti-rabbit IgG secondary antibody (P/N: 926-68071, LI-COR). Pharmacological agents were used in the following concentrations and time courses: dexamethasone (Dex) (D2915, Sigma; 5 μM, 24 h), mifepristone/RU486 (S2606, Selleckchem; 10 μM, 1 h prior to Dex), cyclosporin A (C1832, Sigma, 1 μM, 1 h prior to Dex), mito-apocynin (HY-135869, MedChemExpress, 1 μM, 1 h prior to Dex), and mito-TEMPO (SML0737, Sigma, 1 μM, 1 h prior to Dex). Unless otherwise indicated, all other chemicals were purchased from Sigma-Aldrich.

### Total internal reflection fluorescence (TIRF) imaging

Imaging was conducted in DMEM media without phenol red (ThermoFisher Scientific), supplemented with 10% FBS (Atlanta Biological) and Antibiotic–Antimycotic (ThermoFisher Scientific/Invitrogen). N2a cells were imaged on a Nikon Eclipse Ti2 inverted microscope equipped with a modular LAPP illuminator, motorized TIRF module and LUN-F TIRF laser unit, iXON 897 Ultra camera, motorized xy stage with Piezo z-drive, PFS autofocus, and SR HP Apo TIRF 100X/1.49 NA oil immersion objective in a stage-top temperature- and CO2-controlled chamber. 100X objective and lasers were regularly maintained and calibrated for TIRF imaging by Nikon field support personnel. Glass bottom dishes were secured for TIRF on the motorized stage by magnetic sample holders, with the objective collar adjusted to 0.17 coverglass thickness and 37°C. Images were obtained in NIS-Elements software with 512 × 512 resolution. Cells were located using mCherry signal by epifluorescence microscopy. Cells with flat morphology were chosen to observe maximum cell surface area in the evanescent wave, and thereby compatibility with TIRF. TIRF field and laser angle were set manually for each cell in the GFP channel, to maximize signal clarity and minimize shadow, unless otherwise specified. For experiments using mCh-CD63-pHluorin, videos were acquired of a single field of view and exclusively in the GFP channel to minimize microscope and Piezo calibration time between successive images. Images were acquired with minimum laser power (1-3%) every second for 3 minutes (180-181 total frames) to observe release events while limiting photobleaching and phototoxicity. Following video acquisition, a 2-channel image of the chosen field of view was acquired using GFP and DSRED lasers. For experiments using CD63-pHluorin in combination with mCherry subcellular markers, TIRF fields and angles were set manually for both GFP and DSRED lasers, and 2-channel images were acquired every second for 3 minutes (180-181 total frames per channel). For all experiments, dishes were maintained at 37°C and 5% CO2 during image acquisition. 4-6 fields of view were captured per dish before visualizing a subsequent dish.

### TIRF imaging analysis

TIRF image processing and analysis were performed using Fiji/ImageJ. Events were manually identified by playing raw videos and counted using the multi-pointer tool and ROI manager. Events were further confirmed by measuring size and visualizing sudden increases in fluorescence intensity using plot z-axis profile. Events that were counted as sEV displayed a sudden increase in pHluorin signal (an increase >1000 AU over 10 seconds or less) that had displacement no greater than 6 pixels (0.96um) following appearance. This quantification represents the appearance of pHluorin signal, inclusive of events with signal that either disappeared or were sustained throughout the 3-minute timecourse of video acquisition. Events that were counted were also assessed for small, punctate size with diameter between 2-6 pixels (0.32-0.96um). Punctate signal of appropriate size but which exhibited any of the following, were excluded: oscillating movement following appearance, large displacement following appearance, flickering movement in and out of the field of view. pHluorin signals with the appropriate characteristics but over the size limit were counted only if 2 or more overlapping appearance events were distinctly identifiable. Where indicated, some images are displayed as maximum z-projections of full-length videos and have been processed to reduce background. These images are created by generating the rolling average over 5 frames based on an imageJ macro available at: https://imagej.net/ij/macros/Stack_Moving_Average.txt. Next, the median frame of the video is subtracted from each frame of the video using the image calculator tool to reduce background signal from structures present throughout the duration of acquisition. Finally, the resulting z-stack is converted into a maximum projection. Still frames from the background-subtracted video are used as indicated. All other TIRF data are included as videos of the 5-frame rolling average with no additional background reduction. Videos are compressed to jpeg format, display 20 frames per second, and attached as supplemental files with .avi format.

### Immunocytochemistry

Fixed N2a cells were incubated overnight at 4°C with the following primary antibodies: EEA1 rabbit monoclonal antibody (1:200), Rab7 mouse monoclonal antibody (1:500), Rab27a rabbit monoclonal antibody (1:50), HGS/ Hrs rabbit polyclonal antibody (1:1000), CHMP-4b rabbit polyclonal antibody (1:500), or TOM20 rabbit monoclonal antibody (1:200). Coverslips were washed and subsequently incubated for 1 h at room temperature with the following secondary antibodies: Alexa Fluor 647 goat anti-rabbit IgG highly cross-adsorbed secondary antibody (A-21245; ThermoFisher Scientific/ Invitrogen) and Alexa Fluor 647 goat anti-mouse IgG highly cross-adsorbed secondary antibody (A-21235; ThermoFisher Scientific/ Invitrogen). Coverslips were stained for 10 min at room temperature with 3uM DAPI and dried overnight in the dark. Coverslips were mounted in VECTASHIELD Antifade mounting medium (H-1000, Vector Laboratories) and sealed with clear nail polish.

### Immunofluorescence super-resolution imaging

Images were acquired on the Zeiss LSM 800 confocal microscope running Zen2 software. For HyPerRed-Mito analyses, images of 10 random fields of view were acquired at the z-position with highest HyPerRed-Mito signal intensity using a 40X objective and 2048 x 2048 resolution. For colocalization experiments, cells expressing GFP-nSMase2 were identified and 1.5-3.5um z-stacks were acquired at 0.13um intervals using 63X objective and 2100 x 2100 resolution by Airyscan. Airyscan images were processed by default 3D image settings using the Zen2 software. Further image processing and analysis was performed using Fiji/ImageJ software.

### Colocalization and fluorescence intensity analysis

In Fiji/ImageJ, multi-channel images were split into component fluorescent channels. Z-stack channels corresponding to subcellular marker staining (EEA1, Rab7, or Rab27a) were thresholded using the Yen method and converted into binary masks. The JaCoP plugin was used to calculate Mander’s colocalization coefficients, M1 and M2, between each mask and the z-stack of GFP-nSMase2 signal from the same multi-channel image. Mander’s M1 value, which corresponds to the percent of the mask (for mCh-CD63, EEA1, Rab7, or Rab27a) also occupied by GFP-nSMase2, is reported. To measure fluorescence intensity, z-stack images were collapsed by SUM z-projection and individual cells and nuclei were manually segmented using the polygon selection tool. Multi-channel images were split and the channel corresponding to HyPerRed-mito signal was then used to measure the mean fluorescence intensity of each cell.

### Neutral sphingomyelinase activity assay

N2a cells were seeded at a density of 300-400,000/ well in 6-well plate format and transfected 24 h later with 3ug pEGFP-nSMase2 N1 using Lipofectamine 2000 (ThermoFisher Scientific) according to manufacturer’s instructions. 24 h post-transfection, cells were treated with vehicle, 5uM Dex, or one of the drug paradigms listed above (see “Antibodies and pharmacological agents”) for 24 h. Cells were washed in 1X PBS, transferred to freezer tubes, pelleted by brief centrifugation, and snap frozen in liquid nitrogen, before storing overnight at -80°C. Snap frozen cell pellets were then resuspended on ice in sample buffer (100mM Tris-HCl pH 7.4, 1mM EDTA, 100mM sucrose, 100uM PMSF, 1X Halt protease inhibitor (ThermoFisher Scientific)). Samples were cycled 3 times through 30 seconds of vortexing at maximum speed, 1 minute sonication in a 40W bath ultrasonicator, and 2 minutes incubation on ice. Samples were then centrifuged for 2 minutes at 10,000 rcf, upon which the supernatant containing neutral sphingomyelinase is saved. Activity of the sample is determined by Amplex Red sphingomyelinase assay (ThermoFisher Scientific), according to manufacturer’s instructions. This assay indirectly measures sphingomyelinase activity through an enzyme-couple reaction that converts phosphorylcholine, a by-product of sphingomyelin hydrolysis, into ceramide and finally the fluorescent Resorufin molecule. Reactions were stopped using Amplex™ Red/UltraRed Stop Reagent (ThermoFisher Scientific) and fluorescence measured using the SpectraMax iD5 96-well plate reader with excitation at 530nm and emission at 590nm. nSMase2 enzyme activity was calculated from a standard curve and background fluorescence, measured from siRNA-mediated nSMase2 knockdown, subtracted from samples where indicated. Neutral sphingomyelinase activity was normalized to ug protein in the lysate of each sample.

### Western blot

To collect lysate, cells were collected from plates and lysed in a buffer comprising 150mM NaCl, 25mM Trus-HCl, and 1% Triton X-100. Cell lysate was denaturized by boiling for 10 min at 100°C in 6X Laemmli SDS sample buffer. Proteins were separated by SDS-PAGE on Novex 4-20% Novex tris-glycine gels (ThermoFisher Scientific/ Invitrogen) in Tris-Glycine/SDS buffer (Bio-RAD) and then transferred to nitrocellulose membranes at 0.3 Amp in cold Tris-Glycine buffer (Bio-RAD). Membranes were blocked in 5% bovine serum albumin prepared in 1X PBST (0.5% Tween20) and incubated overnight at 4°C with the following primary antibody concentrations: tubulin alpha mouse monoclonal antibody (1:5000), GFP rabbit polyclonal antibody (1:5000), mCh rabbit polyclonal antibody (1:2500), Rab27a rabbit monoclonal antibody (1:500). Membranes were washed and incubated for 1 h at room temperature with IRDye 800CW goat anti-mouse IgG secondary antibody (P/N: 926-32210, LI-COR) and IRDye 680CW goat anti-rabbit IgG secondary antibody (P/N: 926-68071, LI-COR). Immunoblots were visualized by LI-COR Odessey 9120 near infrared scanner and bands were quantified by Fiij/ ImageJ using rectangle and plot lane tools.

### Real time quantitative PCR

Total RNA was extracted from hippocampal neurons using TRIzol as previously described (DC Rio, et al 2010 in Cold Spring Harbor Protoc). One microgram of RNA was reverse transcribed directly into cDNA using MultiScribe reverse transcriptase (#4311235, ThermoFisher Scientific/ Invitrogen). Real-time PCR was used for quantification of mRNA expression of nSMase2 (Smpd3-FAM, Mm00491359_m1; ThermoFisher Scientific/ Invitrogen) and GAPDH-FAM (Mm99999915_g1; ThermoFisher Scientific/ Invitrogen) using TaqMan gene expression assay universal PCR master mix (4304437, ThermoFisher Scientific) in MicroAmp optical 96-well barcoded plates (ThermoFisher Scientific) on QuantStudio3 Real-Time PCR machine. Data were calculated using the 2−ΔΔCt method, as described by the manufacturer, and expressed as fold increase over the indicated controls (1.0) in each figure.

### sEV isolation and purification for ExoView analysis

Wildtype N2a or hP301L-Tau N2a cells were treated for 24 h with control, 5uM, or 10uM Dexamethasone in HEK media prepared using 10% exosome-depleted FBS. Cell conditioned media was collected and centrifuged at 3220 x g for 20 min to pellet cell debris. Resulting supernatant was concentrated to a volume of 500uL by centrifugation (3220 x g for 10 min) in Amicon Ultra centrifugal filters (100,000 NMWL; Millipore). Extracellular vesicles, 30-300nm in diameter, were isolated from the processed conditioned media by size exclusion chromatography using IZON’s Automatic Fraction Collector (V1) and 35nm qEVoriginal columns (IZON), per the manufacturer’s instructions. Extracellular vesicles were eluted in 500uL filtered, 1X PBS and stored at -80°C before used in subsequent assays.

### ExoView EV-on-a-chip interferometric imaging and fluorescent antibody tetraspanin detection

Microarray chips pre-coated with capture antibodies against mouse tetraspanins CD81, CD9, and IgG controls (UnChained Labs/NanoView Biosciences) were brought to room temperature and pre-scanned to detect background using the ExoView R100 and ExoView Scanner software (UnChained Labs/NanoView Biosciences). Wildtype or hP301Ltau N2a-derived, SEC-purified EV samples were diluted in proprietary incubation buffer (1:100 for wildtype N2a sEVs and 1:50 for P301Ltau N2a sEVs) and incubated overnight (16-18 h) at room temperature on microarray chips, according to the manufacturer’s instructions. To characterize wildtype N2a-derived sEVs, chips were subjected to several rounds of washes in proprietary Solution A and incubated 1 h at room temperature in a mixture of fluorescently conjugated primary antibodies provided by the manufacturer: anti-CD9 CF488a, anti-CD81 CF555, and anti-CD63 CF64 (each prepared 1:500 in cargo blocking buffer; final concentrations 1:1000). Chips were subsequently washed in proprietary Solutions A, B, and deionized water, and dried according to the manufacturer’s instructions. To characterize P301Ltau N2a-derived sEVs, chips were washed and incubated with proprietary Solutions A, C, and D, according to the manufacturer’s instructions for cargo and surface immunofluorescence staining. Chips were subsequently incubated for 1 h at room temperature with fluorescently conjugated primary antibodies provided by the manufacturer, anti-CD9 CF488a and anti-CD63 CF647 (each prepared 1:500 in cargo blocking buffer; final concentrations 1:1000), and with mouse anti-Tau (MAPT) antibody from clone TOMA-1 recognizing oligomeric tau (Millipore Sigma # MABN819) conjugated to Alexa Fluor^TM^ 555 using the Alexa Fluor^TM^ antibody labeling kit (ThermoFisher Scientific/ Invitrogen; Cat No. A88065) according to manufacturer’s instructions. 1.5 uL Alexa Fluor^TM^ 555 conjugated anti-TOMA-1 (prepared 1:200 in cargo blocking buffer; final concentration 1:400) was used to prepare the fluorescent antibody mixture. Chips were washed and dried as before, according to the manufacturer’s instructions. Chips were placed on the ExoView sample chuck and measured by interferometry and fluorescence imaging using ExoView R100 and ExoView Scanner software. Raw particle size, tetraspanin colocalization, and spot montage data were exported from ExoView Analyzer software; the RIgG and HIgG spots served as negative controls, and only fluorescence values exceeding the baseline RIgG and HIgG fluorescence were counted. All solutions, antibodies, microarray chips, and software were prepared and used per the manufacturer’s instructions (UnChained Labs/NanoView Biosciences). Data was collected for 3 biological replicates (3 technical replicates per sample) for sEVs collected from wildtype N2a cells and 6 biological replicates (3 technical replicates per sample) for sEVs collected from P301Ltau N2a cells. Size distributions were generated by collapsing interferometric measurements from all technical and biological replicates for each genotype.

### Statistical analysis

Graphing and statistics were performed using GraphPad Prism (Version 10.2.3). Unpaired, two-tailed t-tests with Welchs’ correction were used for pairwise comparison; 2-way ANOVA with Sidak’s test for multiple comparisons were used when assessing interactions within and between multiple groups; Brown-Forsythe and Welch’s ANOVA tests were used with Dunnett’s T3 corrections for multiple comparisons when comparing more than 2 groups. Data-points and column data are depicted as mean ± SEM as described in corresponding figure legends. Statistical significance was obtained when P < 0.05. Individual P-values are indicated on graphs, and n-values are indicated in the corresponding figure legends.

## Supplemental Material

Supplemental figures 1 and 2 and captions corresponding to supplemental videos 1-10 are provided in a separate file. Supplemental videos 1-10 are provided in a separate .zip file.

## Data Availability

The data that support the findings of this study are available from the corresponding author upon reasonable request.

## Acknowledgements

N2a cells stably transfected with human P301Ltau-EGFP (2N4R) were a kind gift from Professor Juergen Gotz (University of Queensland, Australia) and Professor Ioannis Sotiropoulos (NCSR Demokritos, Greece). We thank Irla Belli for technical consultation for EV-on-a-chip ExoView characterization and TOMA-1 labeling experiments, the Taub Institute Shared Research Core and Kapil Ramachandran for use of Zeiss and Nikon microscopes, and Ioannis Sotiropoulos for helpful discussions. This work was supported by NIH grants R01NS080967, RF1/R01AG069941, R21AG085473, and CUIMC TIGER grant to C.L.W.

## Author Contributions

MR Burke: conceptualization, methodology, investigation, formal analysis, data curation, visualization, and writing—original draft, review, and editing.

CL Waites: conceptualization, methodology, investigation, validation, and writing/editing.

