## Supplemental figures and legends for "Glucocorticoids regulate small extracellular vesicle (sEV) release via activation of nSMase2"

**Supplemental Data**

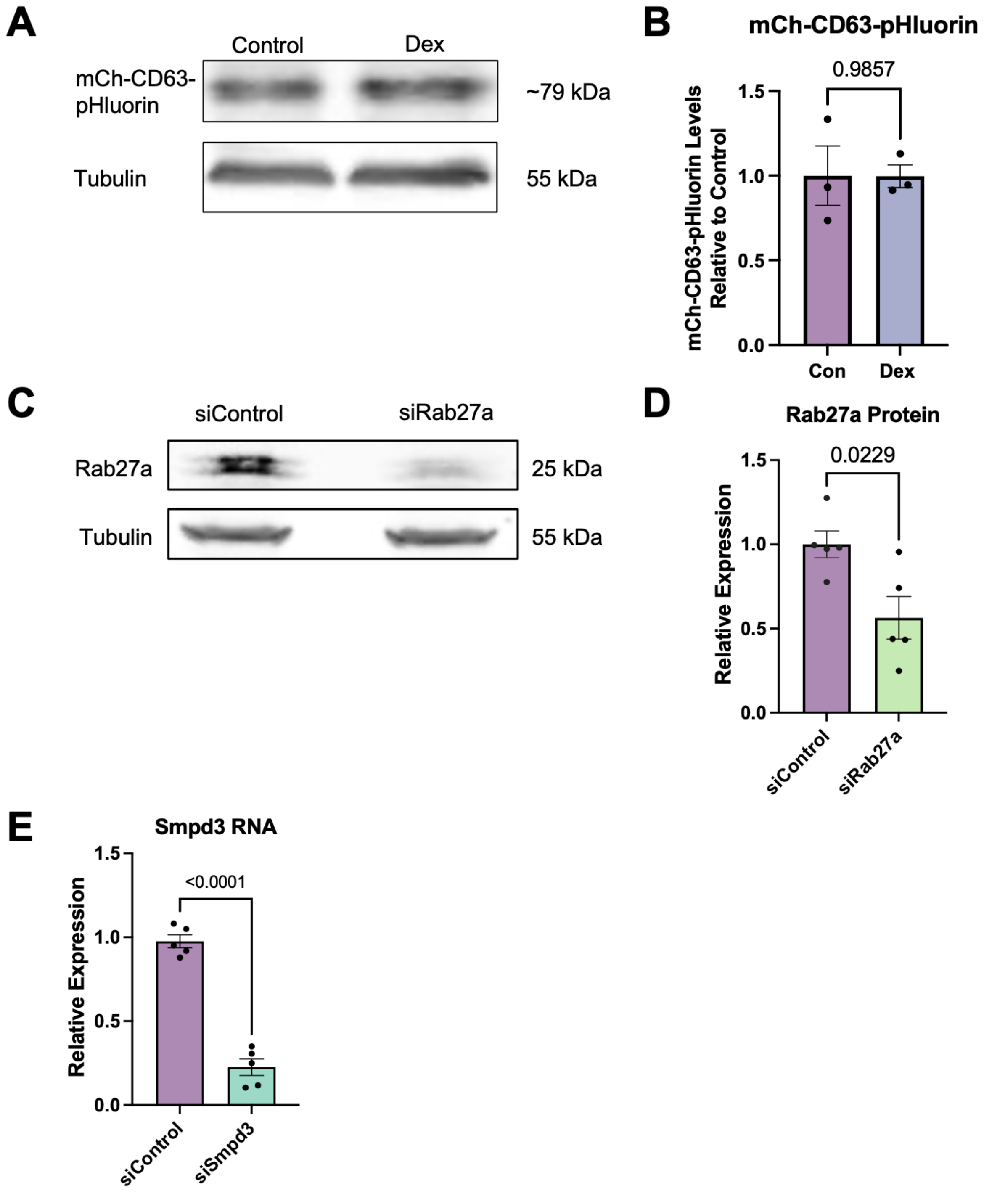

**Figure S1. Immunoblots for effect of Dex on mCh-CD63-pHluorin and knockdown of Rab27a and Smpd3. A-B)** Immunoblot (A) and quantification (B) of mCh-CD63-pHluorin levels in N2a cells following 24 h Dex or vehicle control treatment. The intensity of mCh-CD63-pHluorin (~79kDa) bands were normalized to the average intensity of tubulin (p values shown on graph, unpaired t-test with Welch's correction, n=3). **C-D)** Immunoblot (C) and quantification (D) of Rab27a levels in N2a lysates collected from cells transfected with siRNAs against Rab27a or control for 72 h, normalized to tubulin (p values shown, unpaired t-test with Welch's correction, n=5). **E)** nSMase2/*Smpd3* mRNA levels measured by quantitative PCR, 72 h after addition of siRNAs against either Smpd3 or control (p values shown, unpaired t-test with Welch's correction, n=5). For all graphs, bars represent the mean +/- SEM.

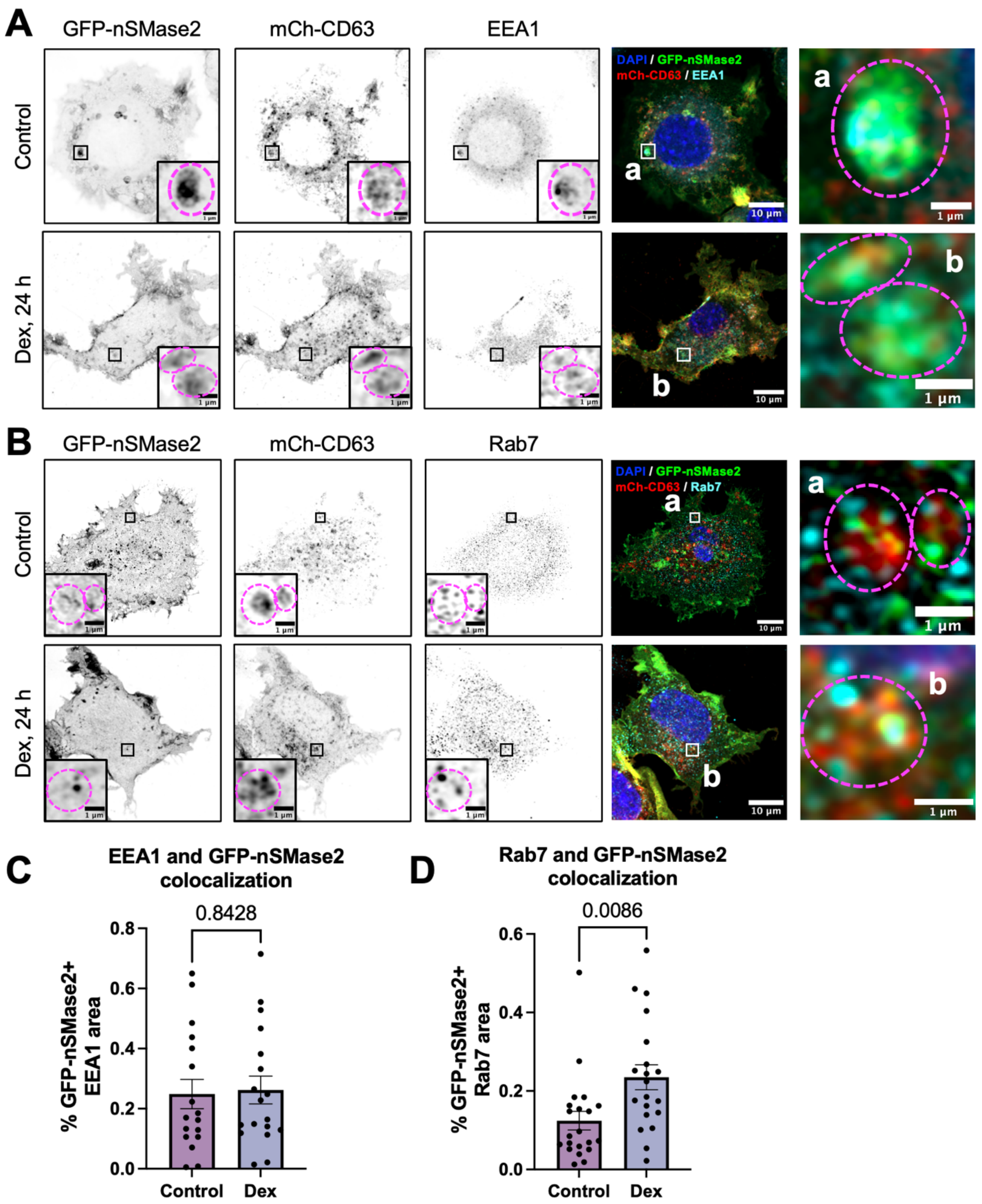


**Figure S2: nSMase2 and CD63 colocalize with early and late endosome markers. A)** Airyscan super-resolution confocal images of N2a cells expressing GFP-nSMase2 (green) and mCh-CD63 (red) and stained for DAPI (blue) and EEA1 (cyan), treated with either vehicle control or Dex for 24 h. Zoomed insets, **a** (Control) and **b** (Dex), are indicated by a black square in black and white images, and white in 4-channel images. Dashed magenta circles indicated the observed area of colocalization of GFP-nSMase2, mCh-CD63, and EEA1. **B)** Airyscan super-resolution confocal images of N2a cells expressing GFP-nSMase2 (green) and mCh-CD63 (red) and stained for DAPI (blue) and Rab7 (cyan), treated either with vehicle or Dex for 24 h. Zoomed insets, **a** (Control) and **b** (Dex), are indicated by a black square in black and white images, and white in 4-channel images. Dashed magenta circles indicated the observed area of colocalization of GFP-nSMase2, mCh-CD63, and Rab7. **C, D)** Quantification of GFP-nSMase2 colocalization with EEA1 (**C**) and Rab7 (**D**), represented as the percent of EEA1 or Rab7 area that is GFP-nSMase2+ as determined by Mander's colocalization coefficient (p values on graph, unpaired t-test with Welch's correction, for EEA1 n=17-18 cells, for Rab7 n= 20-21 cells). For all graphs, bars represent mean +/- SEM.

**Supplemental Video Captions**

**Supplemental Video 1. CD63+ sEV release following 24 h vehicle treatment.** TIRF imaging of mCh-CD63-pHluorin in N2a cells treated with vehicle for 24 h (see **Fig. 1B, 1C (i)**). Images were captured at the TIRF field, magnified by 100X, at 1 frame/second (FPS) for 3 m. Videos were converted into a 5-frame rolling average and exported to play 20 FPS. Scale bar as shown.

**Supplemental Video 2. CD63+ sEV release following 24 h Dex treatment.** TIRF imaging of mCh-CD63-pHluorin in N2a cells treated with 5uM Dex for 24 h (see **Fig. 1B, 1C (ii))**. Images were captured at the TIRF field, magnified by 100X, at 1 FPS for 3 m. Videos were converted into a 5-frame rolling average and exported to play 20 FPS. Scale bar as shown.

**Supplemental Video 3. CD63+ sEV release following 24 h Dex + Mife treatment.** TIRF imaging of mCh-CD63-pHluorin in N2a cells treated with 10uM Mife and 5uM Dex for 24 h (see **Fig. 1B, 1C (iii)**). Images were captured at the TIRF field, magnified by 100X, at 1 FPS for 3 m. Videos were converted into a 5-frame rolling average and exported to play 20 FPS. Scale bar as shown.

**Supplemental Video 4. CD63+ sEV release with siRab27a.** TIRF imaging of mCh-CD63-pHluorin in N2a cells transfected with siRab27a for 72 h and treated with 5uM Dex for 24 h (see **Fig. 2A**). Images were captured at the TIRF field, magnified by 100X, at 1 FPS for 3 m. Videos were converted into a 5-frame rolling average and exported to play 20 FPS. Scale bar as shown.

**Supplemental Video 5. CD63+ sEV release and mCh-Rab27a.** TIRF imaging of CD63-pHluorin and mCh-Rab27a in N2a cells (see **Fig. 2B-D**). Images were captured at the TIRF field, magnified by 100X, at 1 FPS for 3 m. Videos were converted into a 5-frame rolling average and exported to play 20 FPS. Scale bar as shown.

**Supplemental Video 6. CD63+ sEV release and mCh-LAMP.** TIRF imaging of CD63-pHluorin and mCh-LAMP in N2a cells (see **Fig. 2E, 2F**). Images were captured at the TIRF field, magnified by 100X, at 1 FPS for 3 m. Videos were converted into a 5-frame rolling average and exported to play 20 FPS. Scale bar as shown.

**Supplemental Video 7. CD63+ sEV release and RFP-LC3.** TIRF imaging of CD63-pHluorin and RFP-LC3 in N2a cells (see **Fig. 2G, 2H**). Images were captured at the TIRF field, magnified by 100X, at 1 FPS for 3 m. Videos were converted into a 5-frame rolling average and exported to play 20 FPS. Scale bar as shown.

**Supplemental Video 8. CD63+ sEV release with siControl for siSmpd3 following 24 h Dex.** TIRF imaging of mCh-CD63-pHluorin in N2a cells transfected with siControl for 72 h and treated with 5uM Dex for 24 h (see **Fig. 2A, B(i)**). Images were captured at the TIRF field, magnified by 100X, at 1 FPS for 3 m. Videos were converted into a 5-frame rolling average and exported to play 20 FPS. Scale bar as shown.

**Supplemental Video 9. CD63+ sEV release with siSmpd3 following 24 h Dex.** TIRF imaging of mCh-CD63-pHluorin in N2a cells transfected with siSmpd3 for 72 h and treated with 5uM Dex for 24 h (see **Fig. 2A, B(ii)**). Images were captured at the TIRF field, magnified by 100X, at 1 FPS for 3 m. Videos were converted into a 5-frame rolling average and exported to play 20 FPS. Scale bar as shown.

**Supplemental Video 10. CD63+ sEV release following 24 h MitoApo and Dex.** TIRF imaging of mCh-CD63-pHluorin in N2a cells treated with 1uM MitoApo and 5uM Dex for 24 h, representative of other pharmacologic conditions (e.g., CysA+ Dex, MitoTEMPO + Dex) inhibiting mROS (see **Fig. 5D**). Images were captured at the TIRF field, magnified by 100X, at 1 FPS for 3 m. Videos were converted into a 5-frame rolling average and exported to play 20 FPS. Scale bar as shown.
